# From Atoms to Neuronal Spikes: A Multi-Scale Simulation Framework

**DOI:** 10.1101/2025.11.06.686996

**Authors:** Ana Damjanovic, Vincenzo Carnevale, Thorsten Hater, Nauman Sultan, Giulia Rossetti, Sandra Diaz-Pier, Paolo Carloni

## Abstract

Understanding how molecular events in ion channels impact neuronal excitability, as derived from the calculation of the time course of the membrane potentials, can help elucidate the mechanisms of neurological disease-linked mutations and support neuroactive drug design. Here, we propose a multi-scale simulation approach which couples molecular simulations with neuronal simulations to predict the variations in membrane potential and neural spikes. We illustrate this through two examples. First, molecular dynamics simulations predict changes in current and conductance through the AMPAR neuroreceptor when transitioning from the wild-type protein to certain disease-associated variants. The results of these simulations inform morphologically detailed models of cortical pyramidal neurons, which are simulated using the Arbor framework to determine neural spike activity. Based on these multiscale simulations, we suggest that disease associated AMPAR variants may significantly impact neuronal excitability. In the second example, the Arbor model is coupled with coarse-grained Monte Carlo gating simulations of voltage-gated (K^+^ and Na^+^) channels. The predicted current from these ion channels altered the membrane potential and, in turn, the excitation state of the neuron was updated in Arbor. The resulting membrane potential was then fed back into the Monte Carlo simulations of the voltage-gated ion channels, resulting in a bidirectional coupling of current and membrane potential. This allowed the transitions of the states of the ion channels to influence the membrane potentials and vice versa. Our simulations also included the crucial — so far unexplored — effects of the composition of the lipid membrane embedding the ion channels on the membrane potential and revealed a significant impact of temperature on the neuronal excitability. Our combined approaches predicted membrane potentials consistent with electrophysiological recordings and established a multi-scale framework linking the atomistic perturbations to neuronal excitability.

## 1 Introduction

The microscopic changes in the dynamics of an ion channel, due to disease-linked mutations or drugs, profoundly affects its conductance, selectivity, and gating kinetics. These changes alter the inward and/or outward flux of ions through a channel (majorly the Na^+^, K^+^, Ca^2+^, and Cl^−^ ions) which reshapes the action and synaptic potentials of neuronal cells. This dysregulates the excitatory-inhibitory balance, ultimately modifying the excitability of neuronal networks and giving rise to disorders such as epilepsy, schizophrenia, and neurodegeneration. To ultimately link the effects on a molecular level to the brain simulations, one needs to bridge a gap between the different scales. While a significant progress has been made in this respect for neuronal to whole brain simulation^1–5^, the essential link between molecular dynamics - which captures disease-linked molecular alterations - and morphologically realistic neuronal simulations codes, such as NEURON^6^, MOOSE^7^, and Arbor^8^ (developed by some of the authors of this paper, details are found in SI, section I) remains largely unbridged. These codes are based on cable theory^9^, whereas chemical processes are usually modeled using phenomenological approaches, such as Markov schemes or ODEs calibrated to patch-clamp data^10^.

This prospective article provides a proof of concept to couple (i) all-atom molecular dynamics (MD) simulations and (ii) coarse-grained Monte Carlo (MC) simulations with Arbor. (i) The MD studies provide insights on conductance^11^, selectivity^12^, gating transitions^13^, interactions with auxiliary subunits^14^, lipid modulation, allostery, and cooperativity^15^. Simulations at an all-atom level also aid in understanding the conformational transitions^16^ and the effects of small molecules, including binding affinity and energetics^17^, and mutations. Some of these properties may be difficult to access from experiments^18^. In this study, we combined Arbor with MD simulations that compute the single-channel conductance for wild-type and disease-linked variants of α-amino-3-hydroxy-5-methyl-4-isoxazolepropionic acid (AMPA)–type glutamate receptors (AMPARs). This combination allowed us to translate molecular-level alterations into neuronal-scale effects. These cation channels, assembled from GluA1, GluA2, GluA3, GluA4 subunits^19^, mediate fast excitatory postsynaptic currents^20,21^. The GluA2-containing heteromers are the most common AMPAR found in the central nervous system^19,22^. In mature brains, most AMPAR channels incorporate the RNA edited Q607R GluA2 subunit^23^. A cytosine to guanine point mutation in the codon for residue 607 is linked to neurodegenerative diseases and produces the Q607E and R607G variants, which exhibit altered conductance.^24–26^. We performed our MD simulations on a highly similar protein, the rat (r)AMPAR receptor (>98% identical ^27^, to the human (h) AMPAR), for which structural information is available^28^. These all-atom MD simulations were used to estimate the relative change in single-channel conductance caused by each mutation. The corresponding mutation-specific scaling factors were then applied to experimentally measured AMPAR peak conductances in cortical pyramidal neurons^29^ to obtain adjusted conductance values. Arbor then used the adjusted conductances to simulate macroscopic neuronal behavior — specifically, the time courses of membrane potential, *V*_*m*_(*t*), in response to synaptic input. These simulations focused on pyramidal cells - key neurons located in central brain regions, such as the cortex and hippocampus^30^. (ii) The MC simulation code (developed by one the authors, VC) simulates the gating processes in the voltage-gated Na^+^ and K^+^ ion-channels (referred to as VG channels hereafter) ^31^. This code is able to represent the full set of metastable conformational states, despite the coarse-grained nature of the underlying potential of this model. Our simulations included crucial lipid-VG channel interactions^18^ that are absent in standard simulations^31^: indeed, the activation potential of VG channels depends on the surrounding lipid species in a state-dependent manner (that is, on the number of open and closed VG channels^32^). In turn, the (de)activation of VG channels also affect the lipid composition of the neighboring section of the membrane^18^, making the microscopic dynamics of VG channels non-Markovian — namely, dependent on the past depolarization events^33^. This approach predicts the macroscopic currents from the number of open VG channels and also explains the experimental phenomena such as hysteresis and long-term memory effects. Here, the predicted currents carried by K^+^ and Na^+^ ions through VG channels embedded in membrane patches informed Arbor, which calculated the *V*_*m*_(*t*). The latter, in turn, modulated the gating of VG channels, creating a loop between the two.

By integrating the mutation-dependent single-channel conductances from MD simulations into Arbor, and by coupling the latter with MC simulations: we establish a proof-of-concept framework for multi-scale simulations, connecting the changes at atomic-level to neuronal spikes.

## 2 Methods

### 2.1 MD — Arbor simulations

#### MD simulations

Our calculations were based on the rAMPAR open channel cryoEM structure (PDB ID: 5WEO)^28^. This consists of four GluA2 subunits with auxiliary protein Stargazin (TARP γ2). Here, we kept the transmembrane portion of the channel (residues 510-625 and 785-1200). The N-termini were capped with an acetyl group, and the C-termini with a methyl group. The protein was inserted into a POPC lipid bilayer, consisting of 220 lipid molecules in each leaflet. The system was subsequently hydrated with ~48,000 water molecules. To neutralize the system and achieve a salt concentration of 0.30 M, ~260 K^+^ and ~260 Cl^−^ ions were added, for a total of ~223,000 atoms. The dimensions of the pre-equilibrated rectangular simulation box were approximately [150, 150, 120] Å^3^. The position Gln (607) is 586 in rAMPAR, which was mutated to Arg (Q586R), Gly (Q586G) and Glu (Q586E). The wild-type and all mutated systems were generated using CHARMM-GUI^34–36^.

The CHARMM36m force field^37,38^ was used to describe the protein, lipids, and ions, while the TIP3P model^39^ was employed for water molecules. A standard 6-12 Lennard-Jones (LJ) form of the van der Waals potential was used, with force-switched truncation over the range of 10-12 Å. The integration timestep during the production run was 2 fs. The SHAKE constraint method^40^ was applied to chemical bonds involving hydrogen atoms. Constant pressure (1 bar) and temperature (30^°^C) were kept by a Monte Carlo barostat^41^ and a Langevin thermostat with a friction coefficient of 1 ps^−1^, respectively.

The system underwent steepest descent energy minimization for 5,000 steps. Then, velocities for every atom were assigned from a Maxwell-Boltzmann distribution at a temperature of 30^°^C. The system was equilibrated using the CHARMM-GUI proposed six-step protocol^42^, after which the system was further relaxed by 100 ns MD in NPT. Then, NVT MD simulations were carried out for 500 ns. A constant electric field was applied along the axis of the pore of the channel to maintain a voltage (*V*_*m*_) of 600 mV across the membrane^43,44^ during the production run, following Gumbart et al. method^45^. The voltage across the membrane (*V*_*m*_) during the MD simulations was fixed during the entire run. The position of each atom was saved every 10 ps for subsequent analysis. Energy minimization, equilibration and production simulations for all setups were performed using the Amber simulation program^46,47^.

The current (*I*) through the channel was calculated by counting the number of ion crossings using the MDTraj code^48^ (see SI for details, section IV). The conductance was then calculated as *g* = *I/V*_*m*_, where *V*_*m*_ is the voltage across the membrane.

#### Arbor Simulations

We ported the model of a Layer 5b Pyramidal cell described in reference^30^ to Arbor. This provides a data-driven model of phenomena, such as Ca^2+^ spikes and back-propagation of action potentials. Here, (i) we show how the synaptic conductance was calculated, then (ii) we discuss the spatial localization of the stimulus, and finally (iii) the type of synaptic stimulation.

(i) Effective synaptic conductance — the combined contribution of multiple rAMPAR activated at a given synapse — as a function of time was modelled as^49^:

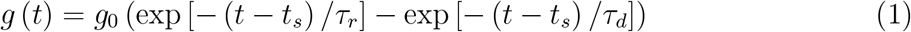

This corresponds to a single incoming spike at time *t*_*s*_, where *g*_0_ = 3.5 nS is the peak conductance, *τ*_*r*_ = 0.5 ms and *τ*_*d*_ = 2 ms are the time constants for rising/decaying flanks. The conductance function in response to multiple spikes is formally the sum of exponentials, but is more commonly implemented as a system of two coupled differential equations. To incorporate the effects of the rAMPAR mutations into Arbor simulations, we considered the experimental effective synaptic rAMPAR conductance, *g*(*t*)^29^, which we approximated and assigned to the most common form of the receptor, Q586R rAMPAR^50^. This is referred to as the baseline form (or baseline species). The approximations which were made here are detailed in the Supporting Information (SI, section V). The effective synaptic conductance for WT and mutants rAMPARs (labeled as MT) then reads as

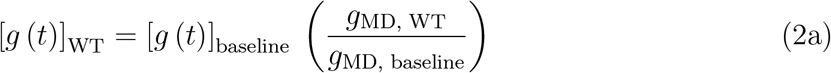

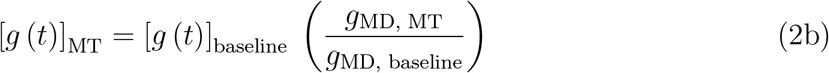

The synaptic current was computed as *I* (*t*) = *g* (*t*) [*V*_*m*_ (*t*) − *E*], where the membrane potential *V*_*m*_ was calculated using the cable equation (initialized as −65 mV) and the synaptic reversal potential of *E* = 0 mV.

(ii) Spatial localization of the stimulus — We consider two types of stimulation towards the cell, representing two extremes in the distribution of inputs along the dendritic tree. Firstly, we considered the localized stimulation, which mimics the experimentally observed patterns of synaptic clustering in vivo, where the co-active excitatory inputs often target the same dendritic branch. Synapses, in this case, were distributed across the dendrite by choosing a random segment of the morphology, and a set of ten random segments in a sphere of 50 µm radius were picked, following the reference^51^. The 50 µm spatial cluster used in our model reflects the biologically observed range of synaptic clustering on dendritic branches, as reported in both experimental and theoretical studies^51–54^. Next, we considered the spatially distributed stimulation, as neurons in vivo often receive distributed excitatory input from a wide array of sources. Here, ten synapses were picked randomly across the dendritic tree. The location of synapses stimulated in this study for both these cases (localized and spatially distributed) are shown in figure 2.

**Figure 1.**
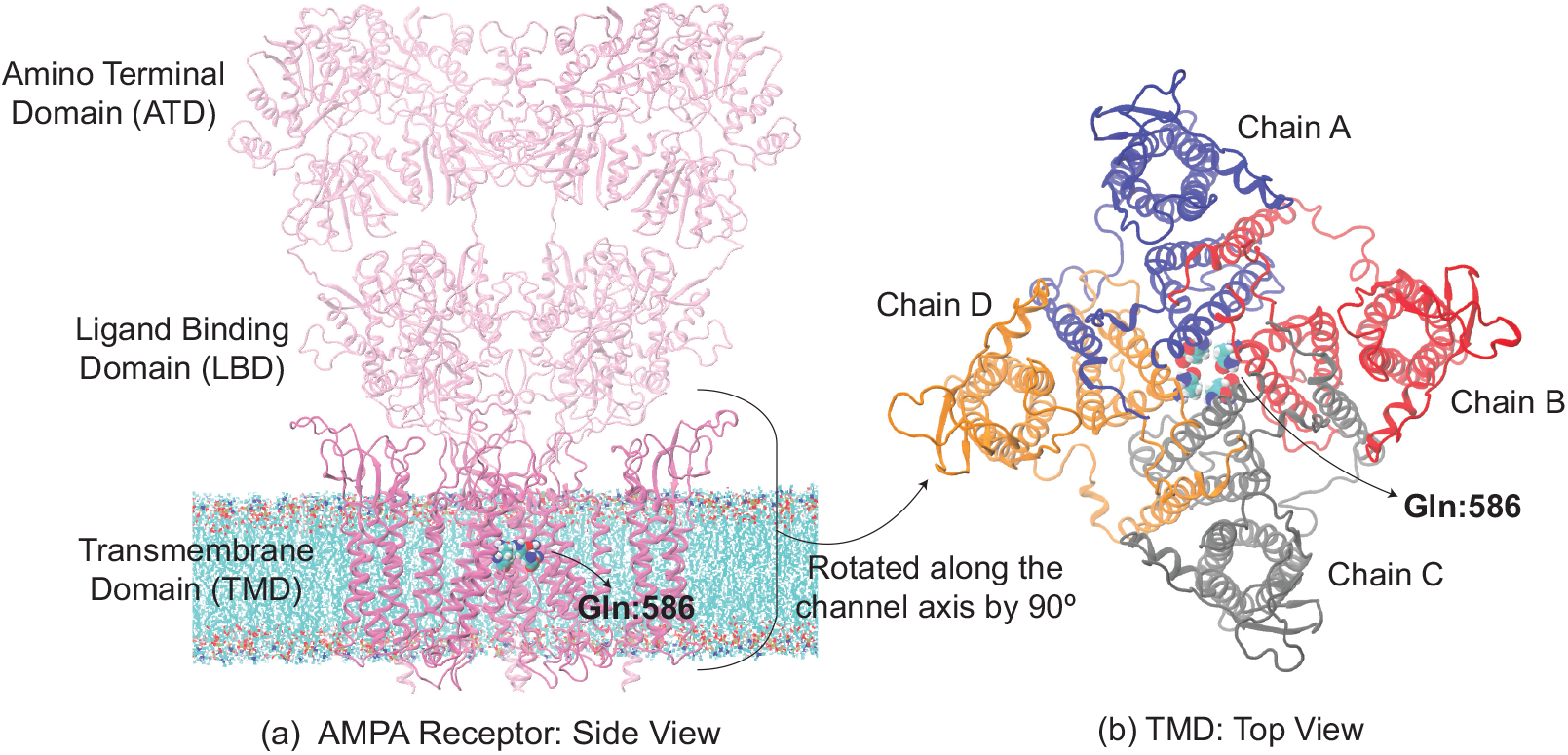
(a) Wild-type rAMPAR (PDB ID: 5WEO) tetramer structure. Gln:586 (in van der Waals spheres representation) corresponds to Gln:607 in hAMPAR. (a) The amino terminal and ligand binding domains in the extracellular region are shown as transparent representations, while transmembrane domain (TMD) as opaque ribbons. The MD simulations are performed on the latter domain which was embedded in the POPC lipid membrane visible in the figure. We display only three chains of the tetramer for clarity here. (b) Top view of TMD. Its subunits (all four chains) are colored differently.

**Figure 2.**
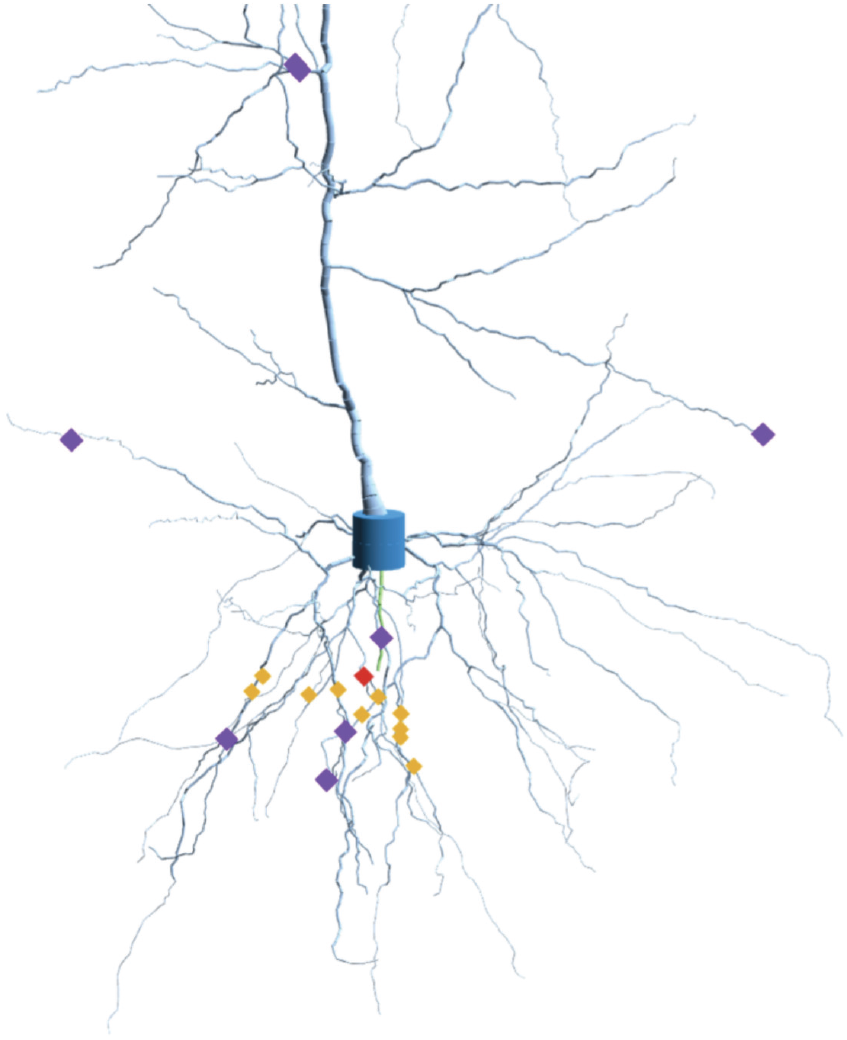
Rendering of the pyramidal neuron used in the study near the soma. This has been generated by producing 3D cylinders of different diameters connected in space describing the reconstructed morphology of a pyramidal cell. The soma has a height of 20 µm and the whole cell roughly fits into a cylinder of height 1300 µm and diameter of 600 µm. Pyramidal neurons are found in the cortex and hippocampus of mammals. The neuron was colored by region: dendrite (light blue), axon (green), and soma (blue). Synapse sites are shown by markers depending on the type of input received: (a) Correlated (orange): picked in a 50 µm sphere around a random center (red). (b) Uncorrelated (violet): Eight of ten random locations used in the control experiment.

(iii) Type of synaptic stimulation — The inputs were produced by individual Poisson point processes (which is the standard way to model spiking activity incoming from other neurons in the network), either correlated or uncorrelated in time. This choice reflects different assumptions regarding the role of pre-synaptic neurons in circuit level function and variability in neural information coding. Injected current was scaled by a synaptic weight; such that, the baseline channel configuration did not elicit spikes under uncorrelated 10 Hz input. The synaptic weight used was 1.5 µS. This provided a threshold baseline to reveal gain-of-function effects under the mutant conditions and to assess the impact of input correlation on neuronal firing. Uncorrelated input was used to define the baseline since it provides the least number of assumptions regarding the functional role of the cell within the network, its location in the brain, and its stage in terms of biological development.

### 2.2 MC — Arbor simulations

#### MC simulations

Our recent MC simulations approaches^33^ are summarized here. A square lattice represented a highly coarse-grained model of a patch of the neuronal membrane. It consisted of VG channels, along with unsaturated/saturated lipids (in grey color in figure 3 (c)). The channels were surrounded by four allosterically linked voltage sensors which allowed the channels to pass from resting to activated states, figure 3 (a,b). The channels were allowed to diffuse and rotate while interacting with the lipids in a state-dependent manner, figure 3 (b). The force field used was an Ising-like potential energy function, which considered “gating” and “interaction” terms. The gating term represents the intrinsic energy differences between the states of the channel (e.g., open vs. closed), while the interaction term accounts for the coupling between the channel and its surrounding lipid environment, such that the energy of a channel state depends on the local lipid configuration. We set the parameters for the simulations as detailed in reference^33^. We performed the MC simulations at three different temperatures: 40^°^C, 30^°^C and 20^°^C. Each simulation was repeated 100 times with different random seeds. The output of the MC simulations was the number of active ion channels and hence the ionic current *I*_*m*_. The latter was the input for the generalized Hodgkin-Huxley model in the Arbor code. We used a simplified cell model: it consists of a soma and a dendrite, each represented by a single cellular volume (also known as compartment). This model was used for simplicity (however, Arbor can also work with more complex cell morphologies). Arbor was then used to calculate the resulting *V*_*m*_(*t*) caused by current clamp stimulations of 80 pA, a value which is within the range used in electrophysiology patch clamp experiments to elicit spiking activity in pyramidal neurons^55^. The resulting *V*_*m*_(*t*) was then the input for the MC simulations, as it modified the gating states (resting and activated) of VG channels which generate *I*_*m*_. The updated *I*_*m*_ was, then again, the input for Arbor, and so on (see SI, section VII and figure II, for details).

**Figure 3.**
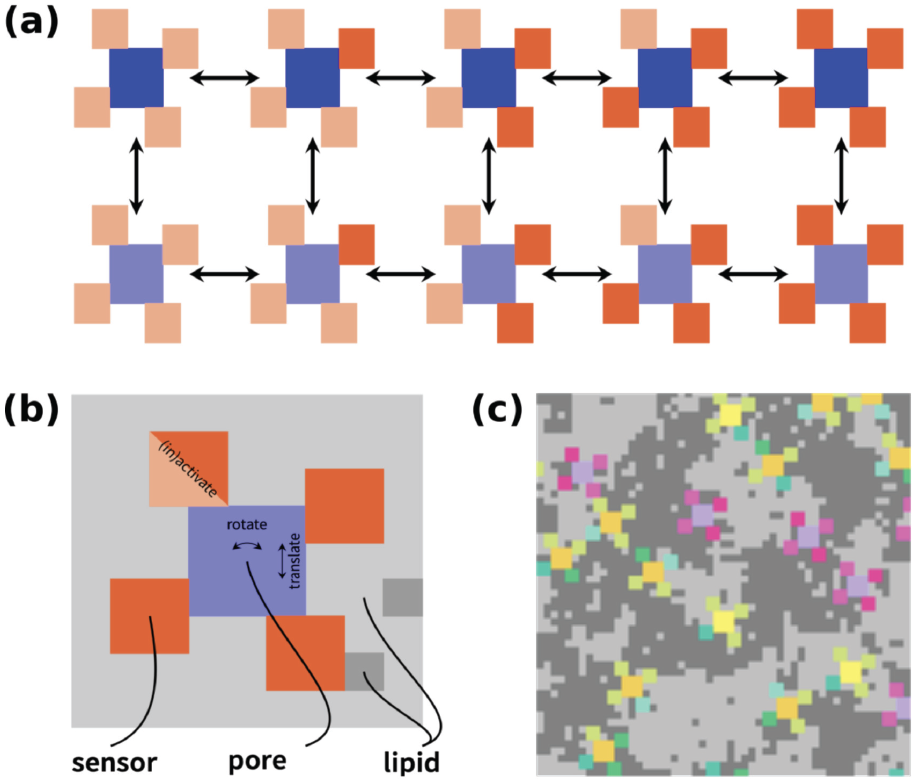
MC simulations. The VG Na^+^ and K^+^ channels (colored in blue in panel (a)) are surrounded by four allosterically linked voltage sensors, in resting (light red color) and activated (red color) states. They can interconvert between the two states by MC steps (arrows on the figure). The channels can rotate and translate within the lipid background, panel (b). Panel (c) shows the channels and lipids are assembled into a square lattice.

## 3 Results and discussion

### 3.1 MD-based calculations

The single-channel conductance, shown in table 1, was calculated as an average over 4 to 7 — 0.5-µs long MD simulations with different initial velocities of the wild-type and mutations: Q586R, R586G, and Q586E rAMPAR tetrameric structures. The results of each individual simulation are displayed in tables I and II of SI, section III. During the mutation studies, each of the four subunits was mutated. For the Q586E mutant, we considered all ionized E586 residues (Q586E.0), 3 ionized (Q586.1) and 2 adjacent ionized residues (Q586.2), while following references^56–58^.

**Table 1:** The MD-averaged conductance values (g_MD_) and their standard errors are reported in the left column. The right column reports the ratio between a specific conductance and that of the baseline (Q586R). Q586E.0-2 correspond to protomer with 0,1, and 2 protons (see text). We report both the actual values and the rounded values used for the neuronal simulations (rounded values are shown in parenthesis).

| rAMPA (system) | Average Conductance ($g_{MD}$) [pS] | Scaling factor |
| --- | --- | --- |
| Wild-Type | 10.2±4.6 | 3.1 (3) |
| Q586R (baseline) | 3.3±1.3 | 1.0 |
| Q586G | 11.7±5.4 | 3.5 (3) |
| Q586E.0 | 92.4±18.3 | 28.0 (30) |
| Q586E.1 | 19.8±6.2 | 6.0 (6) |
| Q586E.2 | 19.8±9.8 | 6.0 (6) |

The MD-derived average conductance of WT rAMPAR (9.6 pS) is within the range of reported experimental values (7-22 pS)^59–61^, and the previous values reported in computational studies^44^. The conductance of Q586R rAMPAR is smaller (3.3 pS), qualitatively consistent with experiment (~0.3 pS)^59,61^, and computational studies^43^. Yet, the conductance of Q586G mutant is instead larger (11.7 pS), indicating that the small, uncharged, glycine preserves functional properties similar to wild-type case of the channel. Our Q586E rAMPAR results may be consistent with the MD studies in reference^26^, which indicate an increase of ion permeability for Q586E rAMPAR. Furthermore, our results for the partially protonated (ionized) Q586E mutants are consistent with the experimentally observed modest increase in conductance in this mutant^25^. This suggests that protonation equilibria at the pore site may dominate under physiological conditions, rather than the fully deprotonated state. We note that our simulations do include the auxiliary subunit Stargazin (TARP *γ*2), which can modify AMPAR conductance^62,63^. Within this limitation, we conclude that the conductance values calculated from our MD simulations for the wild-type and mutant GluA2 receptors are within the range of experimental measurements, validating the ability of our workflow to capture the relevant biophysical changes.

Next, we incorporated these MD calculated conductance ratios into pyramidal neuron simulations by scaling the synaptic conductance parameters of rAMPAR inputs. Specifically, the effective synaptic conductances were obtained by multiplying the experimental value of rAMPAR conductance^29^ by the scaling factor emerging from our MD simulations (table 1). As explained in the methods, we assigned this value to the baseline Q586R rAMPAR which has a scaling factor of 1. From that, we calculated the ratio between current of the mutant and the wild-type and that of Q586R rAMPAR.

From the conductance, we calculated the rAMPAR-mediated single-channel ion currents. Arbor translated them into time courses of membrane potential, *V*_*m*_ (*t*). A spike is elicited when the *V*_*m*_ (*t*) surpasses a threshold of this function, typically between −53 mV and −45 mV in pyramidal neurons^64^. Spikes were calculated by introducing electrical inputs that triggered neuronal responses. We modeled input from ten synapses randomly distributed along the apical dendrites — an arrangement that mimics naturally occurring input patterns, as only a few dozen well-timed inputs could be sufficient to elicit firing, particularly when they arrive close together in space and time^65–67^. Figure 4, together with table III of SI, section VI, presents the *V*_*m*_(*t*) and spike counts for the different rAMPAR variants under both correlated and uncorrelated synaptic input, and the total number of spikes (shown on the top-left corner). The first two rows of figure 4 illustrate localized synaptic stimulation and the last two rows depict spatially distributed stimulation. Simulated time courses of cell membrane potentials turn out to be consistent with electrophysiological recordings of pyramidal neurons in terms of the shape of spikes and frequency (see reference^64^ for a detailed description of the characteristics of spikes in pyramidal cells).

**Figure 4.**
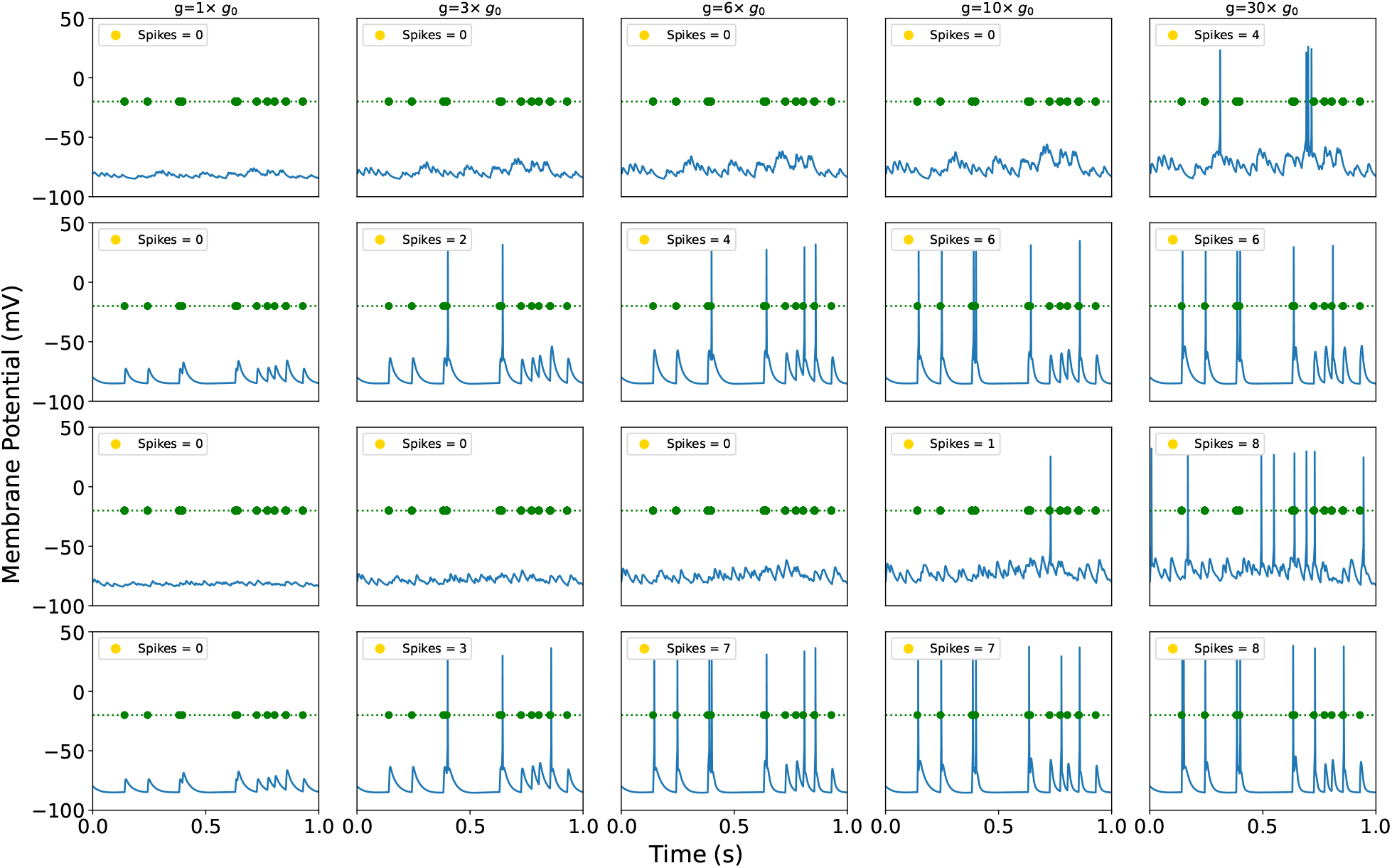
Time series membrane potentials for various AMPAR conductance parameters from MD simulations. Green circle mark input spike times. First and second rows: Localized uncorrelated (first) and correlated (second) synaptic stimulation. Third and fourth rows: Spatially distributed uncorrelated (third) and correlated (fourth) synaptic stimulations.

Figure 4 shows that for both the localized and distributed synaptic stimulation, increasing synaptic strength results in spike formation even when spikes do not occur under baseline conditions (*g* = *g*_0_). This indicates that the mutation lowers the amount of synaptic input required to trigger a spike, thereby increasing the sensitivity of the neuron to stimulation. Under correlated input, a relatively modest increase in AMPAR conductance (*g* = 3 *× g*_0_) is sufficient to elicit spikes, corresponding to the lower end of observed conductance changes and aligning with the wild-type and Q586G GluA2 rAMPAR. As expected, the number of spikes increases with larger values of *g* (figure 4). For uncorrelated input, stronger increase in conductance is required — approximately 10 times *g*_0_ for distributed input and 30 times *g*_0_ for localized input — corresponding to the higher end of conductance modulation (e.g., G586E.0, fully ionized). These results show that the neuron remains more responsive even when input arrives from diverse spatial locations across the dendritic tree (see reference^68^, for example).

In vivo, the number and strength of synaptic inputs required to trigger spikes can vary with dendritic location, neuron type, and synaptic history^69^. While our model explicitly incorporates different protonation states, the effects of other modulatory mechanisms, such as heteromeric subunit assembly, auxiliary protein interaction, and receptor expression levels, are not directly simulated (see SI, section V for a discussion of approximations). However, preliminary modeling of heteromeric AMPAR assemblies suggest conductance levels to be about 10 times *g*_0_, which fall within the range explored here, supporting the physiological relevance of our simulations.

#### MC and Arbor simulations

Our coarse-grain MC simulations predicted the stochastic gating of VG ion channels within a membrane patch.^18^ The MC-based current of K^+^ and Na^+^ ions was given as an input to the Arbor code, which provided the shape and duration of the spikes. The resulting currents differ from that of individual channels, which adhere to Hodgkin-Huxley kinetics^9^. The *V*_*m*_ (*t*) generated from these currents served as an input to the MC simulations, which in turn yield updated current profiles, completing the feedback loop.

The simulations were conducted at different temperatures (40^°^C to 20^°^C). Decreasing the temperature considerably prolonged the *V*_*m*_ (*t*), with values compatible for a variety of neuronal systems^64^, without largely altering the maxima. Inspection of the curves of the spikes (figure 5) leads us to suggest that these temperature-dependent phenomena may arise from this membrane-mediated cooperativity. Indeed, the spike takes much longer to come back down (repolarize) — about 5–6 times longer, on average — while the height of the spike (peak depolarization) stays the same (figure 5). A possible explanation for this is the following: the lipids in a phase-separated membrane form dynamic domains that make neighboring channels ‘help’ each other stay open. Near a phase boundary, small lipid fluctuations can stabilize open states and slow their closure^18^ and the neuronal output may change by altering the lipid mixture, consistently with experimental evidence^70,71^.

**Figure 5.**
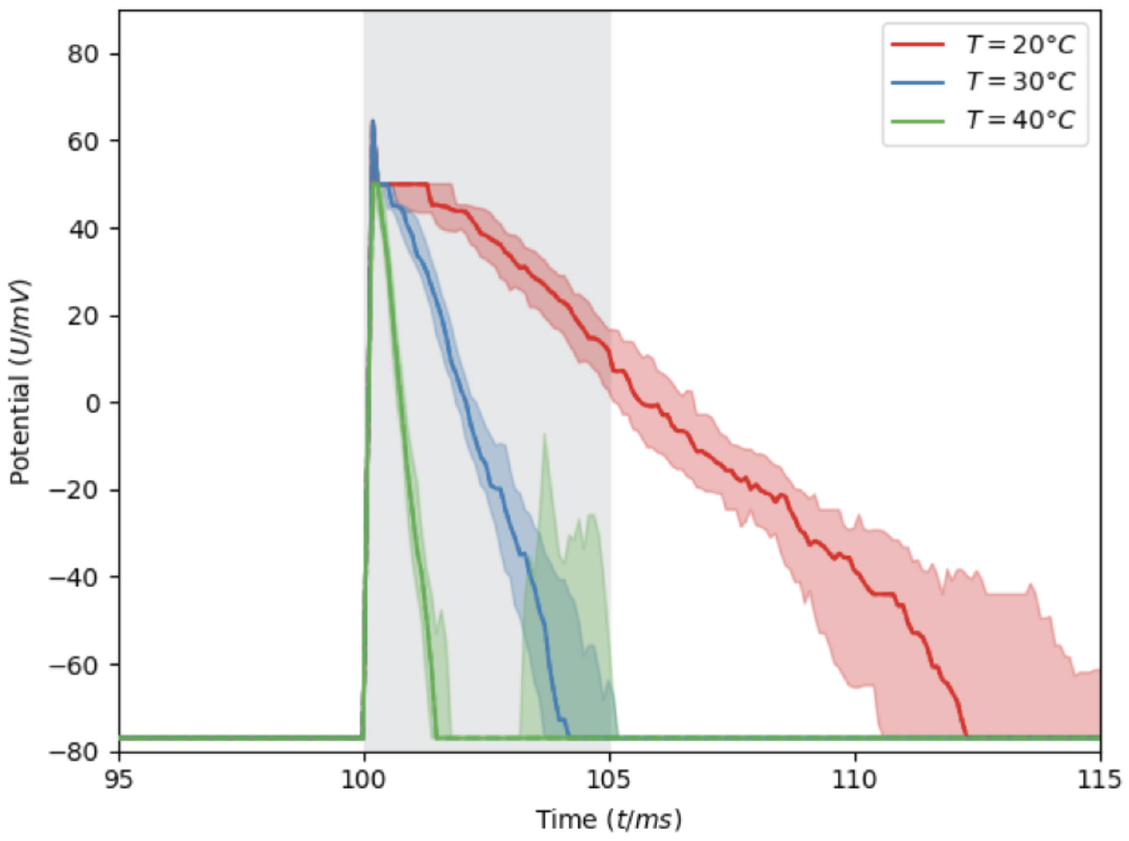
Membrane potential *V*_*m*_(*t*) at three temperature values under specific current clamp stimulations (grey shaded vertical stripe, see SI for details). Shown are the median and area between the first and third quartiles over the 100 MC simulations performed here.

## 4 Conclusions and Outlook

We have presented a framework to directly link molecular and neuronal simulations, thereby bridging a scale that has remained largely unconnected. Although we are still at an embryonal phase and current implementations have limitations (detailed in SI, section V), our results demonstrate that MD- and MC-Arbor couplings provide insight into the mechanistic processes by which molecular changes alter neuronal membrane potentials over time in response to electric stimuli (figures 4 and 5) and this, in turn, affects neuronal excitability. The simulated time courses of cell membrane potentials are consistent with electrophysiological recordings^64^ and demonstrate that our simulations lead to biologically realistic outputs in both cases.

The MD-Arbor scheme could be used to observe changes in neuronal membrane potential, not only for channel variants, but also to investigate the effect of ligand binding, provided that experimental values are available for the system without the ligand. Interestingly, a related MD based approach coupled with a Hodgkin–Huxley model (Chen et al., personal communication) focused on the role of sodium channel selectivity, and how it can shape the time course of membrane potentials. The present framework focuses on synaptic receptor channels, where molecular changes affect excitability through complex temporal integration within morphologically realistic neuronal simulations. While the current implementation is with Arbor, it is expected that the scheme can be implemented with other neuronal modelling software (such as NEURON) with relative ease.

The MC-Arbor coupling introduces a feedback loop by considering how ion channel current in a membrane patch leads to the membrane potential time course and how this modifies ionic current. Our scheme incorporates crucial lipid-channel interactions, which influence the stochastic gating of voltage-gated channels in neuronal membranes.

The scope of these multi-scale approaches could be further expanded by improving the molecular and neuronal approaches presented here. On the molecular side, combining all-atom MD with MC schemes could expand the scope further; for instance, parameters derived from all-atom MD simulations of ion channel activation within the full membrane could inform MC models. This would enable us to investigate the effect of perturbations such as ligand binding, lipid composition or temperature shifts on the excitability of ion channels by bridging time and length scales.

Incorporating important neurobiological processes such as synaptic plasticity and calcium dynamics in neuronal simulations would allow to capture adaptive neuronal behaviour and key aspects of intracellular signalling. Our framework can be readily extended to large-scale neuronal networks, enabling the connection of molecular-level changes to functional outcomes at the network level and potentially to emergent, brain-wide activity patterns. This might enable in silico predictions of the effects of drugs on brain activity.

## Acknowledgement

We would like to thank Dr. Han Lu (FZJ/JSC) for the discussions and contributions to the development of Arbor and its multi-scale scientific use cases. A.D. thanks Ed Twomey (School of Medicine, JHU) for useful discussions about the AMPAR. N.S. would like to acknowledge the support of Dr. Bernard Brooks (NHLBI, NIH). MD simulations for this work were ran on LoBoS at (NHLBI) NIH and Rockfish at Johns Hopkins supercomputering clusters. We would also like to acknowledge the contributions from Daniel Sigg for the Arbor/MC code development. Arbor and MCMC simulations carried out on the JSC infrastructure (JUWELS/JURECA).

## Author Contributions

- AD — Co-designed the project, conceptualized and coordinated the MD-Arbor framework, designed and supervised the MD AMPAR component, and wrote the manuscript.
- VC — Ran the MC simulations, co-designed the project and wrote the manuscript.
- TH — Co-designed the project, implemented the interfaces from MD and MC to Arbor, performed the Arbor simulations, produced visualizations and analysis of the results, and wrote the manuscript.
- NS — Conducted the WT and mutations MD simulations of AMPAR, performed analysis of MD simulations and wrote the manuscript.
- GR — Co-designed the project and wrote the manuscript.
- SDP — Co-designed the project, supervised the tool integration, and wrote the manuscript.
- PC — Co-designed the project and wrote the manuscript.

