## Supporting Information for "From Atoms to Neuronal Spikes: A Multi-Scale Simulation Framework"

**Author Affiliations:**
**Ana Damjanovic**: Department of Biophysics and Department of Physics & Astronomy, Johns Hopkins University, Baltimore, MD 21218, USA; National Heart, Lung, and Blood Institute, National Institutes of Health, USA. (Equally contributed, *Email:*)
**Vincenzo Carnevale**: Institute for Computational Molecular Science and Institute for Genomics and Evolutionary Medicine, Temple University, Philadelphia, PA 19122, USA. (Equally contributed, *Email:*)
**Thorsten Hater**: Simulation and Data Lab Neuroscience, Forschungszentrum Jülich GmbH, Jülich Supercomputing Centre (JSC), 52428 Jülich, Germany. (Equally contributed, *Email:*)
**Nauman Sultan**: Department of Physics & Astronomy, Johns Hopkins University, Baltimore, MD 21218, USA; National Heart, Lung, and Blood Institute, National Institutes of Health, USA. (Equally contributed)
**Giulia Rossetti**: Computational Biomedicine, Institute for Neuroscience and Medicine INM-9, Forschungszentrum Jülich GmbH, 52428 Jülich, Germany; Simulation and Data Lab Biology, Jülich Supercomputing Centre (JSC), 52428 Jülich, Germany; Department of Neurology, University Hospital Aachen, Pauwelsstraße 30, 52074 Aachen, Germany.

### I The Arbor Simulator

Arbor[1, 2] is an open-source library for building simulations of biophysically detailed neuron models based on the cable model to describe electrical behavior of individual cells. Cells can be tied into large scale networks, using action potentials or spikes to transmit information over synapses. This overall set of capabilities allows Arbor to model neuronal networks at a level of resolution beyond point models to explore phenomena like dendritic computation. It provides an alternative to software like NEURON[3], but with a strong emphasis on modern hardware, including GPU accelerators, and scalability to large-scale systems. Arbor is written in C++, though most users interface with it through an intuitive, high-level Python interface built on top of the lower-level implementation.

The electrical dynamics in the cable model are driven by charge transport between the compartments of the discretized dendritic tree — described via the cable equation itself — and the exchange of ions across the membrane, typically called ion channels or mechanisms. These are part of the users' models and either given as a system of differential equations to be solved by Arbor internally or by providing currents directly, interacting with the internals of the library through a versioned interface. For example, this interface is used to facilitate live co-simulation of the Ising-membrane model described below through a network connection.

### II RMSD of the Proteins During the MD Simulations

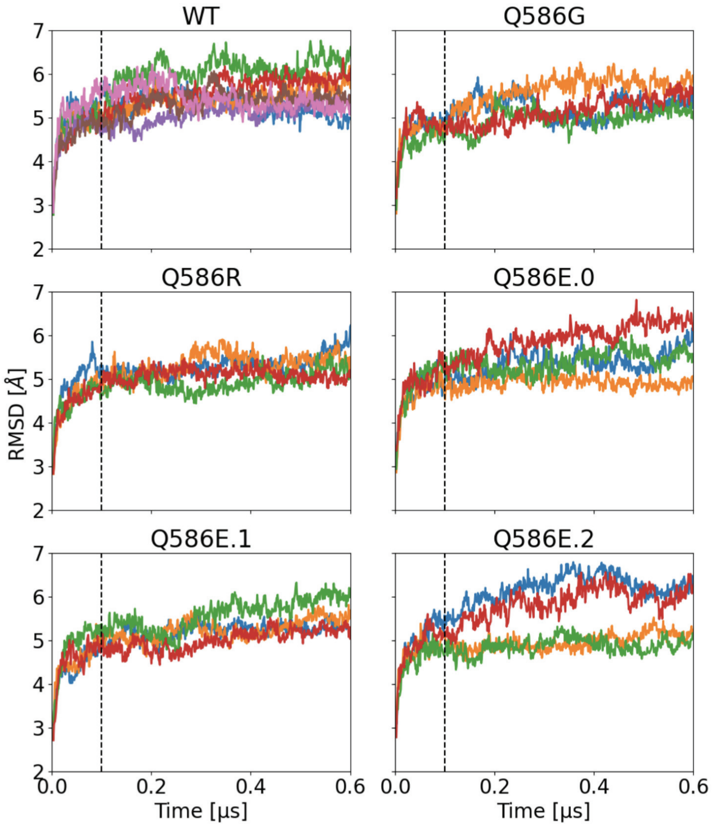


Figure I: Root-mean-square displacement (RMSD) of the protein during the MD simulations. Each panel displays a different mutation, with individual simulation runs color-coded within each. The vertical dotted line at 100 ns marks the beginning of the production runs.

### III MD Simulations Results

| Wild Type | Run 1 | Run 2 | Run 3 | Run 4 | Run 5 | Run 6 | Run 7 | Average | Std |
|---|---|---|---|---|---|---|---|---|---|
| $K^+$ [n] | 2.5 | 0 | 4 | 39 | 7 | 6 | 66 | 17.8 | 23.2 |
| $Cl^-$ [n] | 0 | 0 | 0 | 0 | 0 | -9 | 0 | 1.3 | 3.2 |
| Cond [pS] | 1.3 | 0.0 | 2.1 | 20.8 | 3.7 | 8.0 | 35.2 | 10.2 | 12.2 |

Table I: The net number of potassium and chloride events and conductance calculated during the production runs for WT AMPAR.

| | Run 1 | Run 2 | Run 3 | Run 4 | Average | Std |
|---|---|---|---|---|---|---|
| Q586R | | | | | | |
| $K^+$ [n] | 6 | 0 | 3 | 8.5 | 4.4 | 3.2 |
| $Cl^-$ [n] | -7 | 0 | 0 | 0 | 1.75 | 3.0 |
| Cond [pS] | 6.9 | 0 | 1.6 | 4.5 | 3.3 | 2.7 |
| Q586G | | | | | | |
| $K^+$ [n] | 37 | 1.5 | 46 | 3.5 | 22.0 | 19.9 |
| $Cl^-$ [n] | 0 | 1 | 0 | 0 | 0.25 | 0.43 |
| Cond [pS] | 19.8 | 0.3 | 24.8 | 1.9 | 11.7 | 10.8 |
| Q586E.0 | | | | | | |
| $K^+$ [n] | 162 | 185.5 | 267 | 74.5 | 172.3 | 68.6 |
| $Cl^-$ [n] | -2 | -1 | 0 | 0 | -0.75 | 0.8 |
| Cond [pS] | 87.6 | 99.6 | 142.6 | 39.8 | 92.4 | 36.6 |
| Q586E.1 | | | | | | |
| $K^+$ [n] | 20 | 71 | 18 | 33.5 | 25.6 | 21.3 |
| $Cl^-$ [n] | -1 | -5 | 0 | 0 | -1.5 | 2.1 |
| Cond [pS] | 11.2 | 40.6 | 9.6 | 17.9 | 19.8 | 12.4 |
| Q586E.2 | | | | | | |
| $K^+$ [n] | 95 | 38.5 | 3 | 10 | 36.6 | 36.2 |
| $Cl^-$ [n] | -1 | -1 | 0 | 0 | -0.5 | 0.5 |
| Cond [pS] | 51.3 | 21.1 | 1.6 | 5.3 | 19.8 | 19.6 |

Table II: The net number of potassium and chloride events and conductance calculated during the production runs of mutation study.

### IV Ion Current Calculations from MD Simulations

We used MDTraj[4] to calculate the ion-crossings and calculated the current from these crossings, based on the following references[5, 6, 7]. Crossings were defined as ions traversing an imaginary cylindrical region with a radius of 20 Å and a length designated by the average positions of residues Gln (residue 586) and Asp (residue 590). A complete crossing was recorded if an ion entered one perpendicular end of the cylinder and exited the other. A half-event occurred if an ion either entered the cylinder and remained within the region at the end of the simulation, or if an ion was already within the region at the start of the production run, which were also accounted for in this analysis. The direction of ion crossing defined the sign of the current density. For $K^+$ ions, current was considered positive if they moved along the direction of the applied electric field and negative if they moved in the opposite direction. Conversely, for $Cl^-$ ions, these signs were reversed. The current for simulation time, $t$, was calculated as $I = (n_c - n_a)\,e/t$, where $e$ is the elementary charge, $n_c$ is the net number of cation events, and $n_a$ is the net number of anion events, both measured along the direction of the applied electric field.

### V Assumptions and limitations in simulation

#### MD-Arbor simulations

For our proof-of-concept simulations, we make several simplifying assumptions:

1. Channel gating: We assume the Q/R site mutations do not alter the basic gating kinetics of AMPARs. This is plausible because the mutations are located in the pore rather than in the gating machinery.

2. Calcium permeation: The Q/R site mutation is expected to affect calcium permeation, we do not consider it at this point

3. Synaptic plasticity: We do not model plasticity mechanisms that could change the number or distribution of AMPARs at synapses over time.

4. Subunit composition: We focus on homomeric GluA2 receptors for which the open channel structure is available, even though most physiological AMPARs are heteromeric assemblies of GluA1-4 subunits.

5. Mutation representation: We assume that all AMPAR channels carry the mutation, while in patients heterozygous for GRIA2 mutations (which encodes the AMPA receptor subunit GluA2), only a subset of receptors would actually contain mutant subunits.

Future work will address assumptions 2-5 to better approximate physiological conditions.

#### MC-Arbor simulations

Arbor simulations currently omit the presence of chemical changes (neuroepigenetic processes, for instance), synaptic plasticity, calcium signaling, and long-term changes in dendritic structure or receptor trafficking. Within these limitations, the predicted significant changes in spiking behavior without invoking plasticity underscores the potency of single-channel properties in shaping neuronal output, as seen before[8, 9, 10].

Our MC–Arbor interface modeled the membrane as parallel neuronal patches, each containing two voltage-gated ion-channel types (sodium and potassium, and potassium selective voltage-gated channels) embedded in a binary lipid mixture. This simple configuration serves as a proof of concept: by simulating many patches in parallel, the framework naturally captures disorder and fluctuations (no two patches are identical) and history dependence (each patch's membrane state encodes past activation). However, Arbor could use much more complex cell morphologies as an input.

The MC scheme lacked lipidomes that include sphingomyelin, cholesterol (established ion channel modulators[11]). The implementation of those is underway.

In spite of the several limitations and assumptions used here, the consistency between our calculations and experimental results at different scales, from molecular to neuronal, does support our multi-scale simulations.

### VI Single Cell Simulation Results

| Spatial | Temporal | $g/g_0$ | Spikes |
|---|---|---|---|
| Distributed | Uncorrelated | 1 | 0 |
| Distributed | Uncorrelated | 3 | 0 |
| Distributed | Uncorrelated | 6 | 0 |
| Distributed | Uncorrelated | 10 | 1 |
| Distributed | Uncorrelated | 30 | 8 |
| Distributed | Correlated | 1 | 0 |
| Distributed | Correlated | 3 | 3 |
| Distributed | Correlated | 6 | 7 |
| Distributed | Correlated | 10 | 7 |
| Distributed | Correlated | 30 | 8 |
| Localized | Uncorrelated | 1 | 0 |
| Localized | Uncorrelated | 3 | 0 |
| Localized | Uncorrelated | 6 | 0 |
| Localized | Uncorrelated | 10 | 0 |
| Localized | Uncorrelated | 30 | 4 |
| Localized | Correlated | 1 | 0 |
| Localized | Correlated | 3 | 2 |
| Localized | Correlated | 6 | 4 |
| Localized | Correlated | 10 | 6 |
| Localized | Correlated | 30 | 6 |

Table III: Spike counts with respect to spatial and temporal stimulation pattern and synaptic strength.

### VII MC Simulations and Arbor — Description of the Coupling Interface

The MC model is coupled to a simulation of a single cell in Arbor via a two-way adaptor which enables information exchange between both simulations (see figure II). The adaptor comprises two parts: First, driving the MC simulation for a time step, receiving the membrane potential from Arbor, and sending back the number of open channel pores for each ion species, and second, an ion channel-like object that computes the ionic currents based on the open pores received and sends the membrane potential to the MC simulation. The connection between both parts is established via a network connection. This choice was made due to software restrictions (license incompatibilities); however, it allows flexible deployments as a secondary benefit: simulations may be split across hardware easily and a single adaptor may drive multiple MC simulations, each representing a small patch of the cell's membrane.

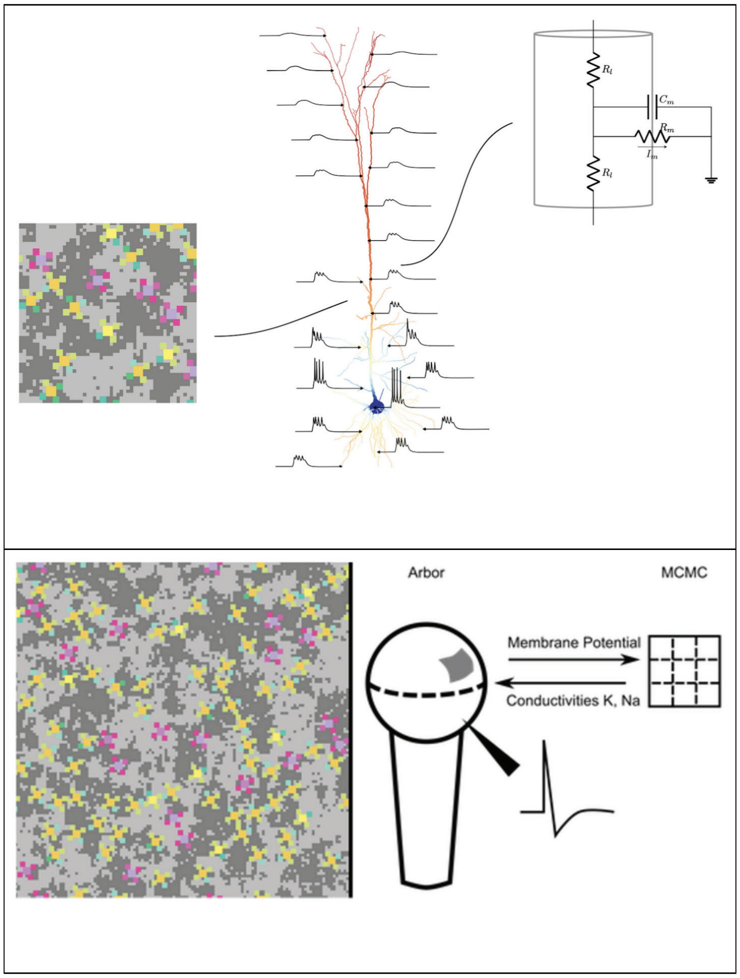


Figure II: Proof of concept of our hybrid MC and Arbor simulation scheme. (Top) Membrane-Cell Co-Simulation diagram, where the membrane potential is calculated by the cable equation and the conductivities of ion channels are calculated by the MC model. (Bottom Left) MC simulation of a membrane patch comprising ion channels and lipids. Shown is a mixture of saturated and unsaturated lipids (grey) interacting with sodium (yellow) and potassium (purple) ion channels on a 128×128 grid. A grid cell corresponds roughly to 1µm$^2$. This patch is embedded into a ball-and-stick cell model in Arbor. (Bottom Right) Setup used in the experiment. Arbor and the external MC software exchange information over a dedicated network socket connection. Arbor provides metadata (timestep, duration) and the membrane potential. The MC simulation replies with the per-ion conductivity derived from the number of open pores.
